# Differential functions of FANCI and FANCD2 ubiquitination stabilize ID2 complex on DNA

**DOI:** 10.1101/2020.02.03.931576

**Authors:** Martin L. Rennie, Kimon Lemonidis, Connor Arkinson, Viduth K. Chaugule, Mairi Clarke, James Streetley, Laura Spagnolo, Helen Walden

**Affiliations:** Institute of Molecular Cell and Systems Biology, College of Medical Veterinary and Life Sciences, University of Glasgow, Glasgow, UK; Scottish Centre for Macromolecular Imaging, University of Glasgow, Glasgow, UK

## Abstract

The Fanconi Anemia (FA) pathway is a dedicated pathway for the repair of DNA interstrand crosslinks, and which is additionally activated in response to other forms of replication stress. A key step in the activation of the FA pathway is the monoubiquitination of each of the two subunits (FANCI and FANCD2) of the ID2 complex on specific lysine residues. However, the molecular function of these modifications has been unknown for nearly two decades. Here we find that ubiquitination of FANCD2 acts to increase ID2’s affinity for double stranded DNA via promoting/stabilizing a large-scale conformational change in the complex, resulting in a secondary “Arm” ID2 interphase encircling DNA. Ubiquitination of FANCI, on the other hand, largely protects the ubiquitin on FANCD2 from USP1/UAF deubiquitination, with key hydrophobic residues of FANCI’s ubiquitin being important for this protection. In effect, both of these post-translational modifications function to stabilise a conformation in which the ID2 complex encircles DNA.

## Introduction

Repair of DNA damage is an important aspect of cellular biology and numerous pathways have evolved to combat different types of DNA damage [1]. Fanconi anemia (FA) is a rare genetic disorder that arises due to mutations within any of the Fanconi Anemia Complementation group (*FANC*) genes, the products of which are involved in repair of DNA interstand crosslinks (ICLs) [2,3] as well as in the maintenance of genomic stability in response to replication stress [4,5]. While quite rare in the general population, FA pathway genes are frequently altered in cancer patients [3].

A key step in this pathway is the ubiquitination of a pair of paralogous proteins, FANCI (~150 kDa) and FANCD2 (~160 kDa) [6,7], which promotes their retention on chromatin [7,8]. In particular ubiquitination of FANCD2 has been shown to be indispensable for cellular resistance to mitomycin C [6,9], which promotes ICLs. Unlike typical ubiquitination events, FANCI and FANCD2 are each specifically monoubiquitinated at a single conserved lysine. Ube2T-FANCL are the E2-E3 pair that mediate ubiquitination [10,11]. In many eukaryotes, including humans, FANCL is incorporated into a pseudodimeric ~1 MDa complex which is known as the FA core complex [12-14]. Removal of the ubiquitins, on FANCI and FANCD2, is also critical for the FA pathway, and this deubiquitination step is catalysed by the USP1-UAF1 complex [15,16].

Evidence suggests FANCI and FANCD2 are involved in recruitment of other proteins [8,17]. However, the mechanistic and structural details of the role of ubiquitination remain ambiguous. The two proteins have been shown to associate *in vivo* [7] and form a heterodimer *in vitro* [18]. A crystal structure of the non-ubiquitinated mouse FANCI-FANCD2 (ID2) complex revealed that each paralog has an extensive α-solenoid fold contorted into a saxophone-like shape [18]. Interestingly, the ubiquitination target lysines are partially buried at the FANCI-FANCD2 interface, which extends throughout the N-terminal half of the proteins. It has been suggested that, the ubiquitin conjugated on FANCI interacts with FANCD2 [19]. The presence of DNA promotes ubiquitination of both isolated FANCI and ID2 complex *in vitro* [20,21], but it is currently unknown how this is achieved. While isolated FANCI and ID2 complex are well known to bind various DNA structures [18,20-23], isolated FANCD2 is less well established to bind DNA [18,22,24]. A FA patient mutation in FANCI, R1285Q, which reduces ubiquitination of the ID2 complex [20,21], has been suggested to reduce both FANCI and ID2 DNA binding, as well as FANCI interaction with FANCD2; however, the magnitude of reduction in DNA binding contrasts between the studies [20-22].

Although FANCD2 monoubiquitination has been documented for almost two decades [6] and FANCI monoubiquitination for over one decade [7], the molecular function of these modifications has been elusive. This has been largely due to the difficulty in isolating pure monoubiquitinated FANCI and FANCD2 proteins for *in vitro* studies. Recent advances in the understanding of the Ube2T allosteric activation by FANCL have allowed for the development of an engineered Ube2T which retains FANCI/FANCD2 lysine specificity but displays enhanced monoubiquitination activity [25]. This engineered Ube2T has facilitated preparation and isolation of highly purified ubiquitinated FANCI and FANCD2 [26]. Here we have used this approach to reconstitute the human ID2 complex in different states of ubiquitination and have characterized DNA-binding for each state. We show that ubiquitination of FANCD2 significantly enhances binding of the ID2 complex to dsDNA, while ubiquitination of FANCI appears to be dispensable for this purpose. CryoEM maps of ubiquitinated FANCD2 in complex with either FANCI, or ubiquitinated FANCI and dsDNA, demonstrate a closure of the ID2 complex via formation of a new protein-protein interface at the C-termini. This interface is apparently disrupted in the FANCI R1285Q pathogenic mutant. We further demonstrate that ubiquitination of FANCI largely protects the ID2 complex from USP1-UAF1 deubiquitination, which likely contributes to the maintenance of ubiquitination-associated ID2-DNA binding enhancement in the cellular context. Therefore, it appears that ubiquitination of FANCI and FANCD2 have separate functions but converge to facilitate and maintain improved ID2-DNA binding.

## Results

### Ubiquitination of FANCD2 enhances ID2-dsDNA binding

In order to explore whether FANCI and FANCD2 ubiquitination impacts ID2-DNA binding, we first purified ubiquitinated FANCI (I_Ub_) and ubiquitinated FANCD2 (D2_Ub_) using our previously established protocol [26]. We then reconstituted the non-ubiquitinated complex (ID2), the complex with ubiquitin only on FANCD2 (ID2_Ub_), and the complex with ubiquitin on both FANCI and FANCD2 (IUbD2Ub). We employed both gel-based Electro-Mobility Shift Assays (EMSAs; ~4 °C) and solution-based Protein Induced Fluorescence Enhancement (PIFE; 22 °C) [27] to determine apparent binding affinities for dsDNA (32 base pair, IRDye700 labelled; Figure 1). The two techniques, although resulting in different measured affinities, both revealed a striking enhancement of DNA binding upon ubiquitination of FANCD2, which ranged between 7-fold (EMSA; Figure 1A) and 10-fold (PIFE; Figure 1B). DNA binding was not detectable under similar concentrations for isolated D2Ub, suggesting that the ubiquitin on FANCD2 is not required for DNA-binding *per se,* but enables a stronger ID2-DNA interaction. Interestingly, the di-monubiquitinated complex (IUbD2Ub) did not result in a detectable change of dsDNA binding compared to ID2_Ub_. These data suggest that FANCD2 ubiquitination serves to either promote or stabilize ID2-DNA binding.

**Figure 1.**
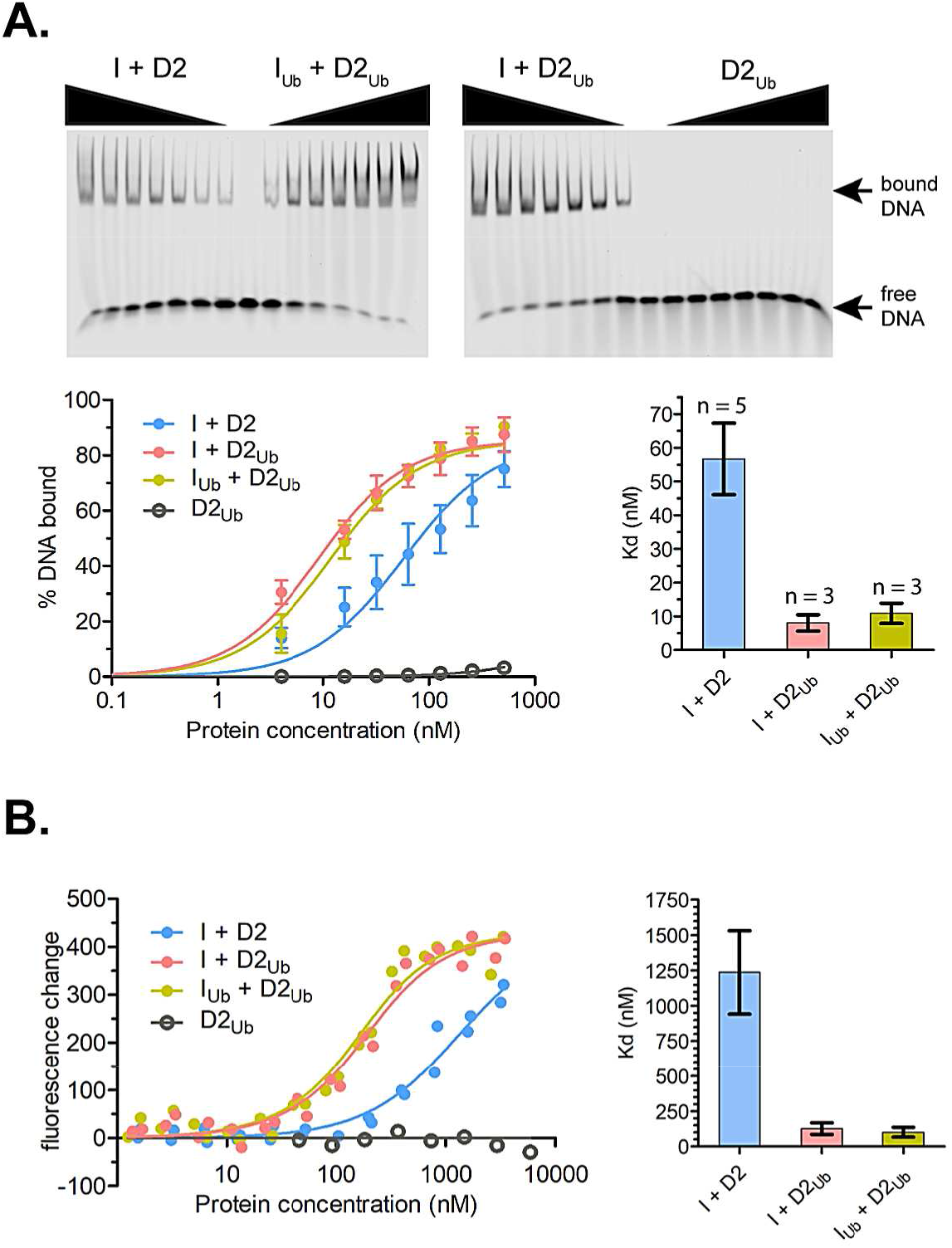
FANCD2 ubiquitination enhances ID2-dsDNA binding. A 32-nucleotide-long IRDye700-labelled dsDNA was used to assess ID2-dsDNA binding, when neither protein is ubiquitinated (I + D2), when only FANCD2 is ubiquitinated (I + D2_Ub_) and when both FANCD2 and FANCI are ubiquitinated (I_Ub_ +D2_Ub_). **A.** *Top*: Infrared signal of free and protein-bound DNA when dsDNA (at 2 nM) was incubated with increasing amounts (4, 16, 32, 64, 128, 256 and 512 nM final) of non/single/double-ubiquitinated His_6_-TEV-V5-FANCI and Flag-FANCD2 protein complexes (I+D2, I_Ub_+D2_Ub_, I+D2_Ub_), mixes were subsequently ran on non-denaturing gels. Ubiquitinated FLAG-FANCD2 at the above concentrations was also assessed for dsDNA binding to examine if FANCD2 has any possible intrinsic DNA-binding affinity upon ubiquitination. *Bottom left*: the percentage of protein-bound DNA to total DNA signal for each ID2 complex concentration (mean ± SD values) is plotted against the various ID2 concentrations, as well as for D2_Ub_ alone. A one-site binding model was fitted to generate binding curves for each of the different ubiquitination states of ID2 used. *Bottom right*: Bar graph showing mean apparent Kd values calculated from the one-site binding model. Error bars: 95% confidence intervals; n: number of replicates. **B.** *Left*: Fluorescence changes of IRDye700 labelled dsDNA (at 125 nM) when incubated at increasing ID2 concentrations (I+D2, I_Ub_+D2_Ub_, I+D2_Ub_ and D2_Ub_), ranging from 1.3 nM to 5.9 μM. Measurement of fluorescence enhancement for each ID2 complex was conducted for two separately prepared complexes and all data points for each protein-combination were used in fitting to a one-site binding model. *Right*: Bar graph showing mean apparent Kd values calculated from the one-site association quadratic equations. Error bars: 95% confidence intervals.

### Ubiquitination of FANCD2 is associated with formation of a secondary ID2 interface

To examine the structural details of enhanced DNA binding affinity we determined cryoEM maps of reconstituted ID2 complexes with ubiquitinated FANCD2, at modest resolutions (Figure 2A, EV1). Reference-free 2D class averages of ID2_Ub_ and I_Ub_D2_Ub_-dsDNA exhibited similar overall shapes, but different to previous non-ubiquitinated ID2 class averages [12], hinting at a gross conformational change upon ubiquitination of FANCD2. Reconstructed maps of ID2_Ub_ and I_Ub_D2_Ub_-dsDNA, at resolutions of 25 and 12 Å, respectively, both exhibited a closed, torus-like shape. Fitting of the mouse truncated ID2 crystal structure into the I_Ub_D2_Ub_-dsDNA map resulted in a poor fit (Figure 2B; *left panel*), but flexible fitting using iMODFIT [28] with secondary structure restraints improved the agreement (cross-correlation score from 0.56 to 0.85). The primary movement occurred for the FANCD2 C-terminal “arm” and resulted in formation of a new interface with the FANCI C-terminal “arm” that closes the ID2 complex (Figure 2B; *right panel*). We refer to this interface henceforth as the Arm ID2 interface. A difference map between the fitted model and the experimental map illustrates tube-like volume, most likely representing the bound DNA. This volume is positioned just below the Arm ID2 interface and encompassed within the torus (Figure 2B; *right panel*), suggesting that formation of the Arm ID2 interface is necessary for this binding conformation. At these resolutions, we are not able to unambiguously place the conjugated ubiquitins. We propose that the closed conformation must be stabilized to tightly bind DNA and ubiquitination of FANCD2 acts for this purpose.

**Figure 2.**
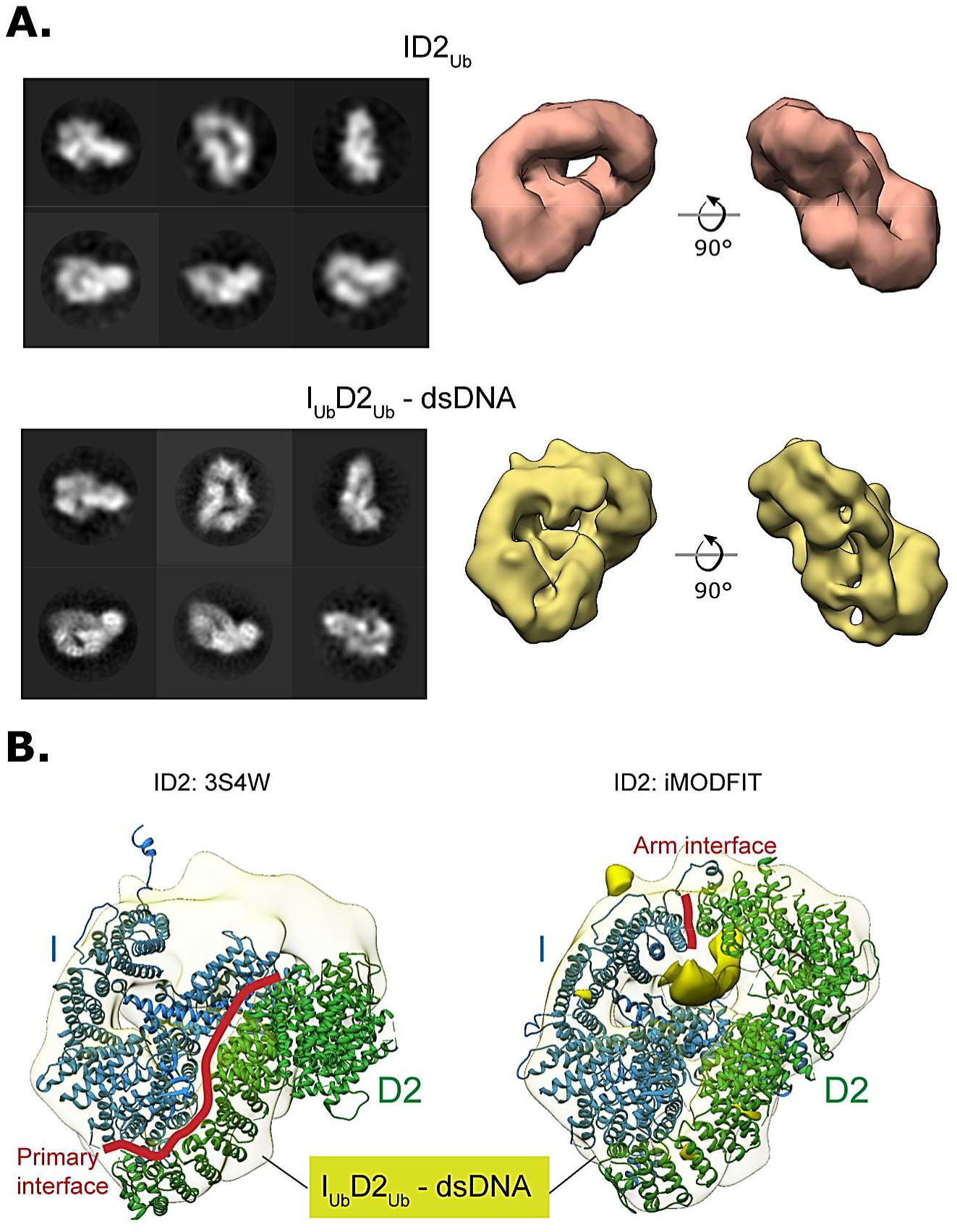
FANCD2 ubiquitination creates a secondary ID2 “Arm” interface, resulting to an altered, closed ID2 conformation. **A.** *Left*: CryoEM 2D classes of ID2_Ub_ and I_Ub_D2_Ub_-dsDNA molecules. *Right*: Resulting 3D reconstructions of ID2_Ub_ and I_Ub_D2_Ub_-DNA. Both structures exhibit a torus-like shape. **B.** *Left*: Rigid body fit of the mouse ID2 atomic structure (3S4W) into the human IUbD2Ub-dsDNA cryoEM map. *Right*: Flexible fit of the mouse ID2 atomic model into the human I_Ub_D2_Ub_-dsDNA cryoEM map (iMODFIT). The C-termini “arms” of FANCD2 and FANCI close in the flexible fitting. A tube-like volume of the IUbD2Ub-DNA cryoEM, just beneath the Arm ID2 interface (solid density), which is not occupied by the ID2 iMODFIT structure, likely represents bound DNA.

### The Arm ID2 interface is required for efficient ID2 ubiquitination

Interestingly, the site of the pathogenic FANCI mutation R1285Q is in proximity to the Arm ID2 interface (Figure 3A). We therefore hypothesized that this mutation may disturb ID2 ubiquitination by preventing the formation of the closed ID2 state. We first examined whether this mutation brings any changes in FANCI’s capacity to interact with DNA and FANCD2, as well as whether it affects its ability to get ubiquitinated. We found that both wild-type (I^WT^) and mutant (I^R1285Q^) proteins could be ubiquitinated *in vitro* to the same extent, and addition of DNA resulted in comparable enhancement of ubiquitination between the two proteins (Figure 3B). Furthermore, by measuring the binding affinities of RED-tris-NTA (Nanotemper) labelled His-tagged I^WT^ and I^R1285Q^ for FANCD2 (using PIFE), we found that the affinities were similar and both in the low nanomolar range (Figure 3C). Nevertheless, the FANCI mutation resulted in an apparent reduction in ID2 ubiquitination (Figure 3D), consistent with previous results [20,21].

**Figure 3.**
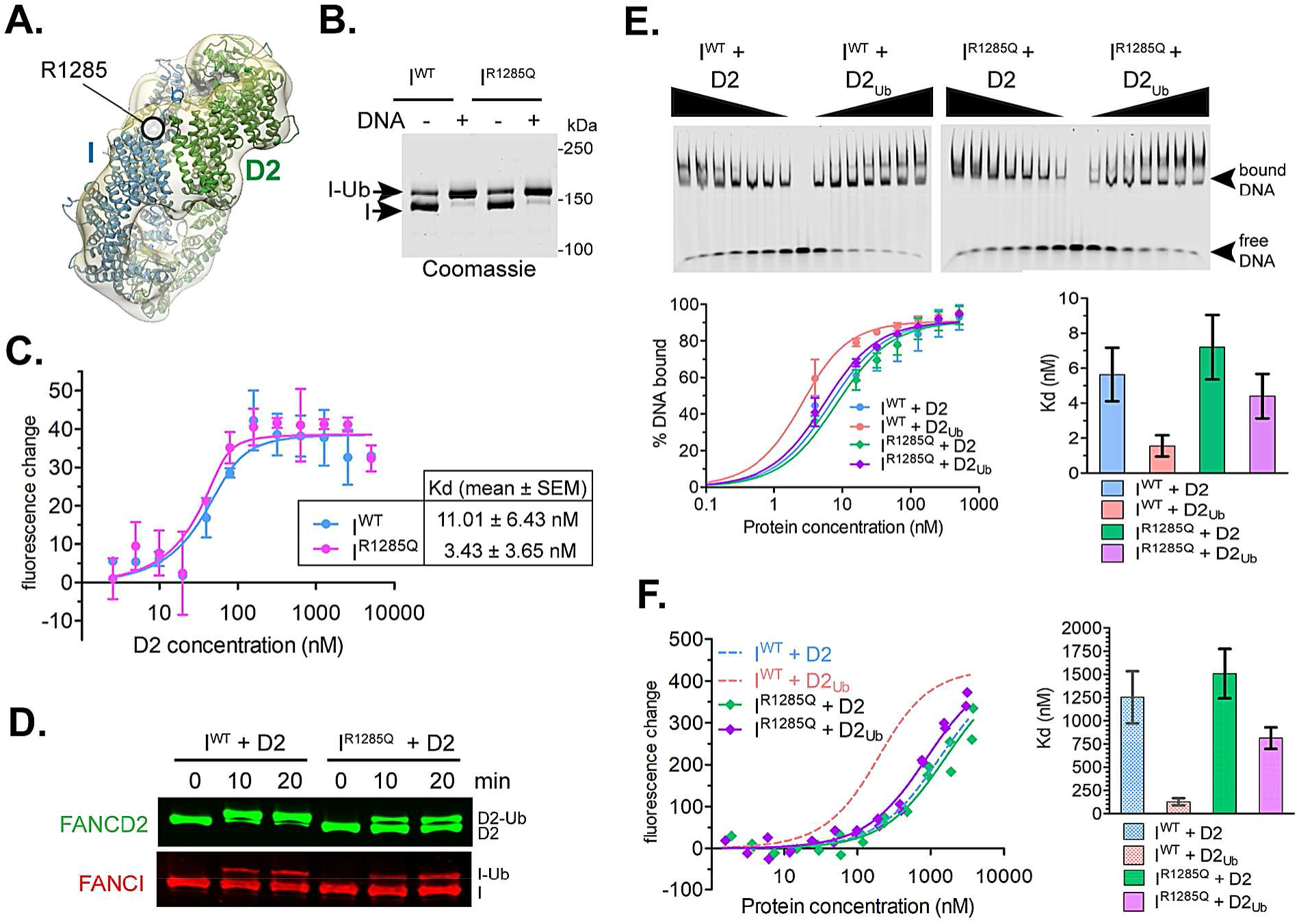
The pathological FANCI R1286Q mutation disrupts ID2 ubiquitination, likely via disturbing the closed ID2 conformation. **A.** Arginine 1285 of FANCI, which is mutated in some FA patients, is located in the Arm ID2 interface in our cryoEM I_Ub_D2_Ub_-dsDNA structure. **B.** Both wild-type (I^WT^) and R1285Q mutant (I^R1285Q^) FANCI can be efficiently ubiquitinated in the presence of DNA. Reactions were performed at 2 μM FANCI substrate for 60 min in the absence or presence of 4 μM ssDNA. **C.** Both wild-type (I^WT^) and R1285Q mutant (I^R1285Q^) FANCI efficiently associate with FANCD2. Fluorescence changes occurring when RED-tris-NTA-labelled FANCI (I^WT^ or I^R1285Q^; both at 60 nM) was incubated at increasing concentrations of FANCD2 (ranging from 2.48 nM to 5.08 μM) were plotted (mean values from two replicates with standard deviation errors-bars shown) for each FANCD2 concentration and apparent Kd values (mean ± SEM) were determined from fitting to a one-site binding model. **D.** FANCD2 within an I^R1285Q^D2 complex is resistant to ubiquitination. Reactions were performed at 4 μM ID2 substrate and 16 μM dsDNA. Progress of FANCD2 and FANCI ubiquitination was monitored by western blotting following SDS-PAGE. **E.** FANCD2 ubiquitination can enhance ID2-DNA binding only when D2_Ub_ is complexed with I^WT^ but not when complexed with I^R1285Q^, as determined by EMSAs. *Top*: Infrared signal of free and protein-bound DNA when IRDye700-labelled dsDNA (at 2 nM) was incubated with increasing amounts (4, 16, 32, 64, 128, 256 and 512 nM final) of His_6_-FANCI and FLAG-FANCD2, and mixes were subsequently ran on nondenaturing gels. *Bottom left*: the percentage of protein-bound DNA to total DNA signal for each ID2 complex concentration (mean values from two replicates with standard deviation error-bars shown) is plotted against the different ID2 concentrations used. A one-site binding model was fitted to generate binding curves for each of the different ubiquitination states of ID2 used. *Bottom right*: Bar graph showing mean apparent Kd values calculated from the one-site binding model. Error bars: 95% confidence intervals. **F.** FANCD2 ubiquitination can enhance ID2-DNA binding only when D2_Ub_ is complexed with I^WT^ but not when complexed with I^R1285Q^, as determined by PIFE. *Left*: Fluorescence changes when IRD700-labelled dsDNA (at 125 nM) is incubated at increasing concentrations of I^R1285Q^ +D2 or I^R1285Q^ +D2_Ub_ (complex concentrations ranging from 1.54 nM to 3.8 μM). Measurement of fluorescence enhancement for each protein combination was conducted for two separately prepared complexes and all data points for each protein-combination were used in fitting to a one-site binding model. *Right*: Bar graph showing mean apparent Kd values calculated from the one-site binding model. Error bars: 95% confidence intervals. I^WT^ + D2 and I^WT^ + D2_Ub_ calculated curves and corresponding Kd range values (Figure 1B) are also shown for comparison.

Interestingly, the reconstituted mutant I^R1285Q^D2_Ub_ complex behaved differently in terms of dsDNA binding, compared to the wild-type ID2_Ub_ complex. EMSAs revealed only a minor reduction of ID2-DNA affinity when FANCI was mutated to FANCI^R1285Q^, and when FANCD2 was ubiquitinated the ID2-DNA affinity was not substantially enhanced, unlike that seen for wild-type complex (Figure 3E). PIFE similarly showed a small increase in I^R1285Q^D2-DNA affinity when FANCD2 was ubiquitinated (Figure 3F), unlike the levels observed with I^WT^D2 versus I^WT^D2_Ub_. Taken together, these results suggest that the FANCI^R1285Q^ patient mutation does not directly alter ID2-dsDNA binding, but instead restricts ID2 ubiquitination and associated DNA binding enhancement. This is likely achieved via disruption of the Arm ID2 interface, seen in the closed ID2 state. Hence, the loss of I^R1285Q^D2 complex ubiquitination can be rationalized if the closed state is important for ubiquitination.

### FANCI ubiquitination protects ID2_Ub_ complex against USP1-UAF1 deubiquitination

The USP1-UAF1 complex specifically targets ubiquitinated FANCD2 for deubiquitination utilizing an N-terminal module of USP1 [29]. Although this can occur when D2_Ub_ is in isolation or in complex with FANCI, di-monoubiquitinated ID2 complexes (I_Ub_D2_Ub_) bound to DNA remain largely resistant to USP1-UAF1 deubiquitination [29,30]. We thus hypothesized that since FANCI ubiquitination is not directly involved in enhancing ID2-DNA binding, its role may be in protecting FANCD2’s ubiquitin from USP1-UAF1 activity. To examine to what extent the presence of I or I_Ub_ influences D2_Ub_ deubiquitination, we compared the progress of D2_Ub_, ID2_Ub_ and I_Ub_D2_Ub_ deubiquitination by USP1-UAF1 (in the presence of dsDNA) in a timecourse (Figure 4A). USP1-UAF1, at low (25 nM) concentrations, deubiquitinated both D2_Ub_ and ID2_Ub_ at similar rates. However, the I_Ub_D2_Ub_ substrate remained almost completely resistant to USP1-UAF1 activity (Figure 4A; *Left*). At higher concentrations of USP1-UAF1 (100 nM), we found that FANCD2 can be deubiquitinated, albeit at slower rate than D2_Ub_ and ID2_Ub_, which are rapidly deubiquitinated in under 10 minutes (Figure 4A; *Right*). These data suggest that access to the ubiquitin on FANCD2 for USP1-UAF1 is reduced when in complex with ubiquitinated FANCI and DNA.

**Figure 4.**
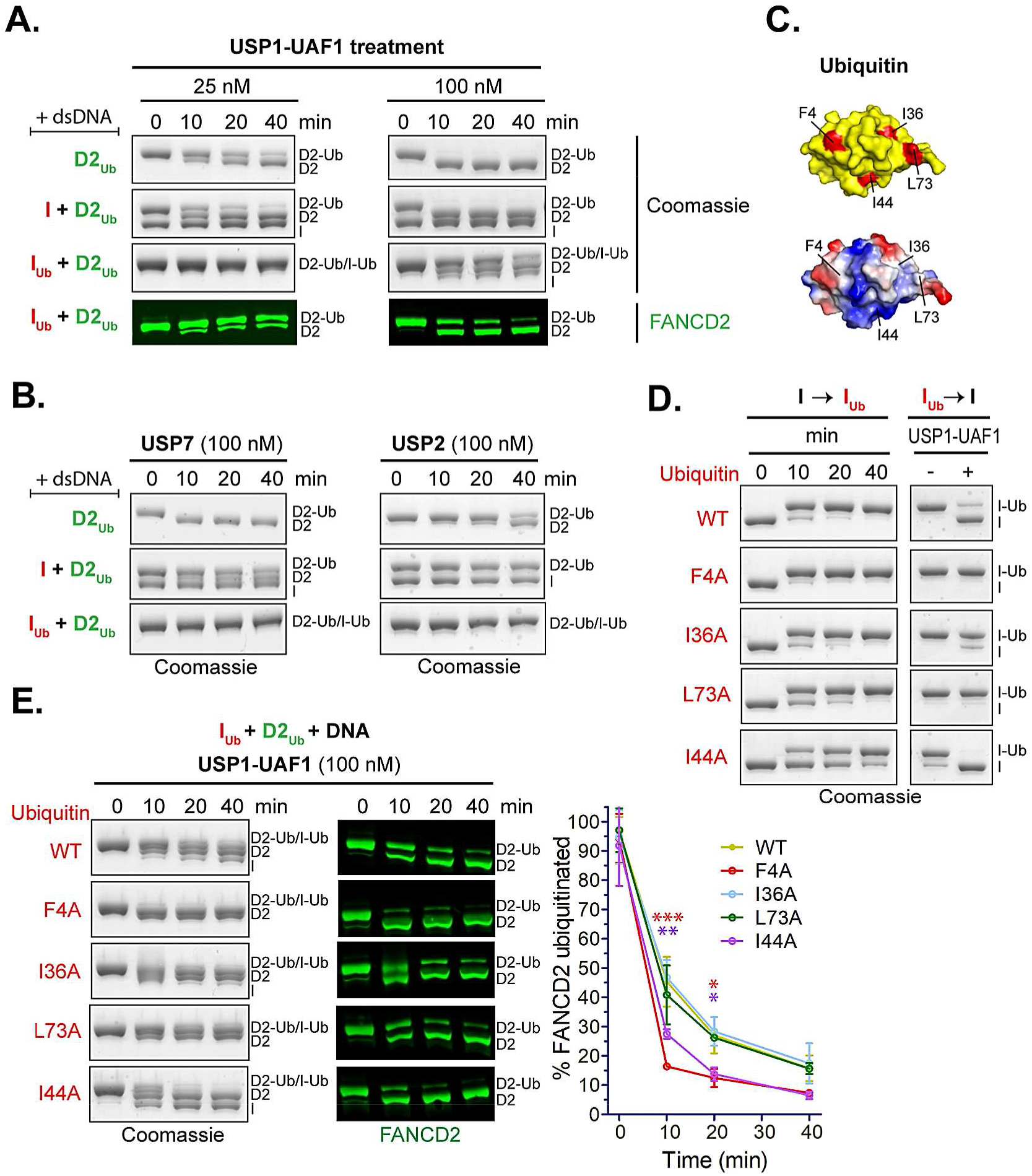
FANCI ubiquitination of ID2_Ub_ complex protects D2_Ub_ from USP1-UAF1 mediated deubiquitination. **A.** USP1-UAF1 can efficiently deubiquitinate D2_Ub_ in the presence or absence of FANCI (I), but not in the presence of ubiquitinated FANCI (IUb). Ubiquitinated His_6_-3C-FANCD2 (D2Ub) was mixed with a 50-nucleotide-long dsDNA and either His_6_-TEV-V5-FANCI (I), ubiquitinated His_6_-TEV-V5-FANCI (IUb) or no protein; protein-DNA mixes were subsequently incubated with USP1-UAF1 (at 25 nM or 100 nM) for indicated time periods. Deubiquitination of D2_Ub_ and I_Ub_ was assessed, following SDS-PAGE, by Coomassie staining of the gels, as well as by western blotting of transferred blots with a specific FANCD2 antibody. **B.** USP7 and USP2 can deubiquitinate D2_Ub_ in isolation, but their activity towards D2_Ub_ is greatly reduced in the presence of FANCI (I) or ubiquitinated FANCI (IUb). Reactions were set-up as in A, but with 100 nM USP7 or USP2. Deubiquitination was assessed, following SDS-PAGE, by Coomassie staining. **C.** Location in the ubiquitin structure (PDB: 1ubq) of the four hydrophobic residues chosen to be mutated to alanine. *Top*: location of F4, I36, I44 and L73 (shown in red) in ubiquitin’s surface (shown in yellow). *Bottom*: same as above but with ubiquitin’s surface coloured according to charge (red: negative, blue: positive, white: no charge). **D.** Time course of FANCI ubiquitination with various ubiquitin mutants and corresponding sensitivity/resistance to USP1-UAF1 mediated deubiquitination of resulting products. Deubiquitination reactions in the presence or absence of USP1-UAF1 (100 nM) occurred for 30 minutes. **E.** Time course of USP1-UAF1 mediated deubiquitination of I_Ub_D2_Ub_-DNA complexes consisting of D2_Ub_ a 50-nucleotide-long dsDNA and I_Ub_ produced with various ubiquitin mutants or wild-type (WT) ubiquitin. Progress of deubiquitination reaction was assessed following SDS-PAGE, by both Coomassie staining of the gels, as well as by western blotting of transferred blots with a specific FANCD2 antibody. FANCI ubiquitination with indicated ubiquitin-mutants (or WT) and subsequent deubiquitination of resulting I_Ub_D2_Ub_-DNA complexes were conducted twice; the residual FANCD2 ubiquitination, calculated from the FANCD2 blots for each time-point, was plotted for each type of ubiquitin in the protein complex (mean ± SD). Statistically significant changes compared to WT ubiquitin (repeated measures ANOVA test with Bonferroni correction) are indicated with asterisks (red: F4A; purple: I44A). * p<0.05, ** p<0.01, *** p<0.001.

Since USP1-UAF1 is a specific deubiquitinase for FANCD2 [15,29], we wanted to assess whether alternative DUBs, that lack FANCD2 specificity (i.e. target structurally diverse substrates), are also able to access the ubiquitin on FANCD2 when this is in complex with I or IUb. We also assayed deubiquitination of D2Ub, ID2_Ub_ and IUbD2_Ub_ by USP7 and USP2 (in the presence of dsDNA) in a time-course (Figure 4B). While USP7 and USP2 could deubiquitinate D2_Ub_ in isolation, the presence of FANCI or ubiquitinated FANCI reduced D2_Ub_ deubiquitination. This suggests that, in the ID2_Ub_-DNA and I_Ub_D2_Ub_-DNA complexes, the ubiquitin on FANCD2 is protected against generic DUB activity. However, only the presence of ubiquitinated FANCI can protect the ubiquitin on FANCD2 from USP1-UAF1 deubiquitination.

One explanation for the requirement of FANCI’s ubiquitin to forming a DUB a resistant complex is that FANCI’s conjugated ubiquitin interacts with FANCD2. Ubiquitin typically interacts with other proteins via any of its three (F4, I36 and I44) hydrophobic patches [31]. Thus, we mutated four key hydrophobic residues of ubiquitin located within these patches, which are at distinct regions of the ubiquitin structure (Figure 4C). Subsequently, we ubiquitinated FANCI (in the presence dsDNA) using either wild-type or any of these four ubiquitin mutants (F4A, I36A, L73A, or I44A). Nearly complete FANCI ubiquitination was achieved for each mutant and wild-type ubiquitin (Figure 4D). We tested each I_Ub_ in DUB assays and found that only I_Ub-I44A_ was also deubiquitinated to the same extent as I_Ub-WT_ by USP1-UAF1. The I36A mutation resulted in partial loss, whereas the F4A and L73A mutations on ubiquitin resulted in complete loss of USP1-UAF1 activity (Figure 4D). To assemble I_Ub_D2_Ub_-DNA complex with ubiquitin mutants on FANCI, we added D2_Ub_ to each ubiquitin-mutated I_Ub_ and subjected the resulting I_Ub_D2_Ub_-DNA complex to deubiquitination treatment with USP1-UAF1 (Figure 4E). Both F4A and I44A ubiquitin mutations, despite having contrasting effects on deubiquitination of FANCI alone, resulted in an increased susceptibility to FANCD2 deubiquitination. The I36A and L73A mutations had negligible effects on FANCD deubiquitination (Figure 4E). The same effect of I44A and F4A on FANCD2 deubiquitination was observed when using four times lower concentrations of USP1-UAF1 in single time-point DUB assays (Figure EV2A). We reasoned that ID2-DNA binding or complex formation were unaffected by these mutations, since WT, F4A and I44A I_Ub_D2_Ub_ complexes were still able to bind DNA efficiently, unlike I_Ub_ alone (Figure EV2B) or D2_Ub_ alone (Figure 1). Taken together, the F4 and I44 hydrophobic surfaces on ubiquitin are important for I_Ub_ mediated protection of D2_Ub_, potentially via interaction with FANCD2.

Interestingly, the ubiquitin on FANCI was also protected from USP2, USP7 and USP1-UAF1 deubiquitination in the I_Ub_D2_Ub_-DNA complex. In fact, in context of IUbD2Ub-DNA complex, FANCI deubiquitination by USP1-UAF1 was even slower than FANCD2 deubiquitination and incomplete by 40 minutes (Figure EV3). However, isolated I_Ub_ was completely deubiquitinated within 30 minutes by USP1-UAF1 (Figure 4D). Taken all together, the sequential action of FANCD2 and FANCI ubiquitination results in a stable I_Ub_D2_Ub_-DNA complex, resistant to deubiquitination.

## Discussion

Monoubiquitination of both FANCI and FANCD2 is a key step in the FA pathway. Here we have demonstrated separate functions for the two monoubiquitination events occurring on the ID2 complex. FANCD2 ubiquitination was found to modulate ID2-dsDNA binding affinity, whereas FANCI ubiquitination was found to largely protect the ubiquitinated ID2 complex from deubiquitination. A simple explanation for our data is that the ID2 complex can explore two different conformational extremes: an open state, as in the previously reported non-ubiquitinated ID2 crystal structure and EM maps [12,18,32], and a closed state, as observed here (Figure 5). We propose that DNA binding allows ID2 to reach the closed ID2 conformation, in which the Arm ID2 interface forms. This conformational change likely exposes FANCD2’s target lysine, as well as its adjacent acidic patch [25] to allow docking of Ube2T and subsequent FANCD2 ubiquitination; the latter modification stabilizes the closed state. However, the action of USP1-UAF1 within the cell will allow a population of ID2_Ub_ to revert to an open state via an unstable closed state of deubiquitinated ID2. The sequential ubiquitination of FANCI may ensure that the majority of ID2 population exists in the closed state, since I_Ub_D2_Ub_ is largely protected from USP1-UAF1 activity. Hence, FANCD2 ubiquitination appears to enhance ID2-DNA binding by reaching the closed state, while FANCI ubiquitination acts to maintain this closed state (and thus the higher DNA affinity), by impairing deubiquitination. As a result, the sequential action of FANCD2 and FANCI ubiquitination is expected to ‘lock’ the ID2 complex on DNA. Since FANCI’s Arginine 1285 is located near the observed Arm ID2 interface in our cryoEM structures, the R1285Q mutation on FANCI is expected to disturb this interface by inhibiting the closed ID2 state and subsequent ubiquitination events that depend on this (Figure 5).

**Figure 5.**
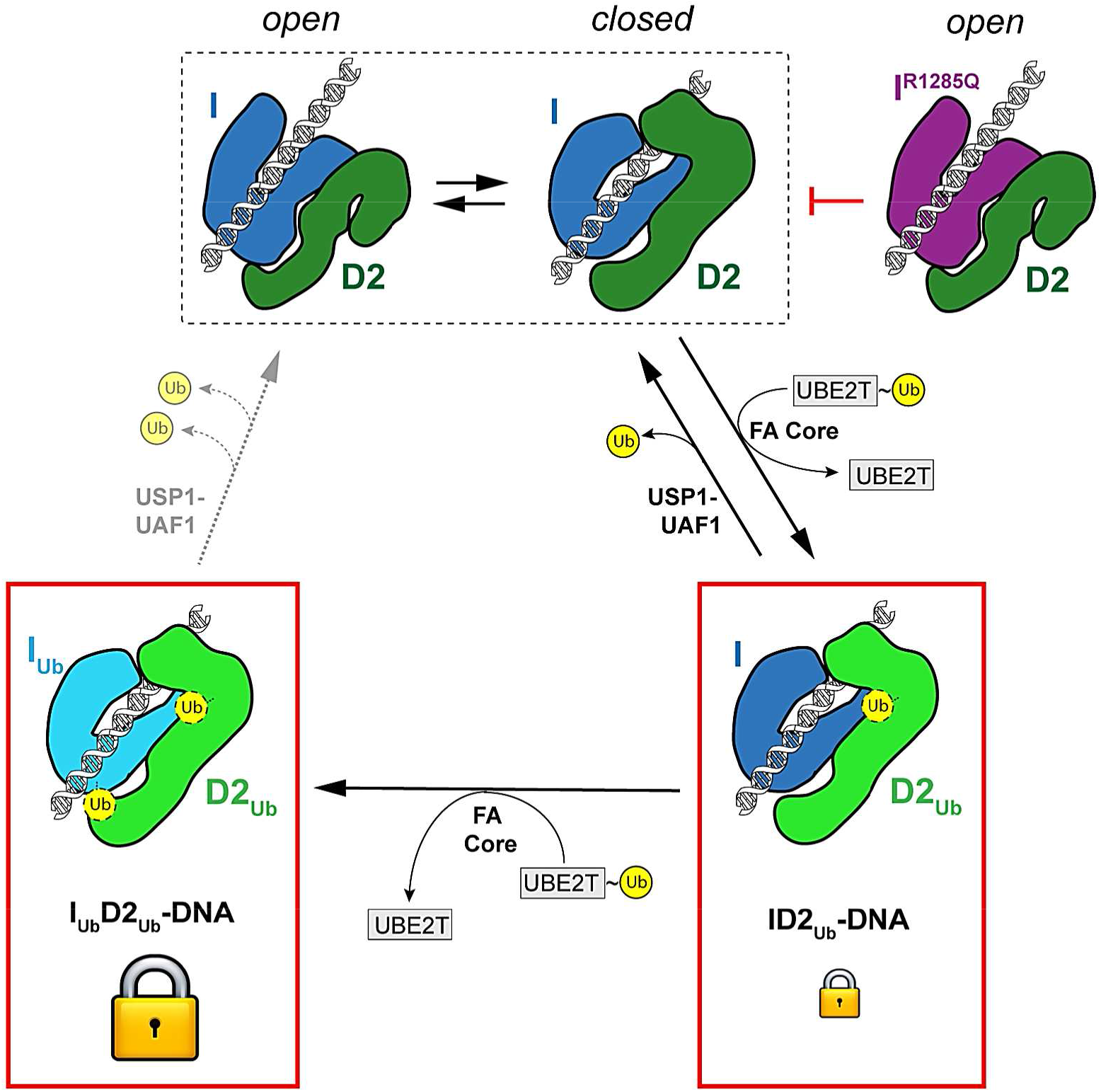
ID2 DNA-dependent di-monoubiquitination may be required for shifting the equilibrium of ID2 complexes to a closed conformation with enhanced DNA binding. Model showing how ID2 interaction with DNA and subsequent FANCD2 and FANCI ubiquitination may enrich for ID2 molecules present in the closed arm conformation: ID2 interaction with DNA is proposed to promote a dynamic equilibrium where ID2 can exist in both an open and a closed conformation. FANCD2 ubiquitination by Ube2T and the FA-core complex, within a closed ID2 conformation status, can shift this equilibrium in favour of the closed conformation, with the action of USP1-UAF1 counteracting such shift. When FANCI is also ubiquitinated by Ube2T and the FA-core complex, both ubiquitins are largely resistant to USP1-UAF1 deubiquitination, and thus the doubly ubiquitinated ID2 can remain in the closed conformation. The latter conformation has a tighter DNA affinity and hence locks ID2 onto DNA. The R1285Q mutation on FANCI likely restricts formation of a closed ID2 conformation, which in turn negatively impacts on ID2 ubiquitination and locking onto DNA.

Here we show that FANCD2 ubiquitination within in the ID2 complex results in enhancement of DNA binding and a conformation that encircles DNA in a closed state. The enhancement of DNA binding and closed state observed upon ubiquitination is supported by three recent studies [33–35], which include cryoEM structures of human I_Ub_D2_Ub_-DNA and chicken ID2_Ub_-DNA [33,34]. Altogether, our data and these recent studies indicate that ubiquitination of FANCD2 is required to form a closed state for ID2 complex. In the I_Ub_D2_Ub_-DNA structure [33], FANCI’s R1285 is indeed located within the Arm ID2 interface, and forms a salt bridge with FANCD2 Q1365. Using our reconstitution approach, we are able to robustly control the ubiquitination status of each protein and determine that the I^R1285Q^D2_Ub_ reduces the enhanced DNA binding observed upon FANCD2 ubiquitination, potentially by reducing the formation of closed state. In addition, we find that FANCI’s ubiquitin does not provide further DNA binding enhancement.

The biological implications of an ubiquitination-dependent locking of ID2 to DNA are currently speculative: this locking may ensure that the ID2 complex is properly recruited to sites of replication arrest where ID2 may be needed. However, it is currently unknown if the locked-on-DNA I_Ub_D2_Ub_ complex is able to slide, recognize a specific DNA structure, or execute another unknown function, which will result in the concentrated ID2 foci frequently observed in nuclei of DNA-damaged cells [7]. Nevertheless, our model is consistent with the observed enrichment of ubiquitinated forms of FAND2/FANCI in chromatin [7,8]. In addition, our model suggests a sequence of events, where a FANCD2-ubiquitination is first required to achieve a shift towards an ID2 state with higher DNA affinity, and then FANCI ubiquitination follows to maintain this state. This is consistent with the observations that FANCI ubiquitination lags behind FANCD2 ubiquitination *in vitro* [25,30].

*In vitro* [25,30] and cell-based [7] assays have shown that the blockage of FANCI ubiquitination also results in reduced FANCD2 ubiquitination. According to our data, this may occur because FANCD2’s ubiquitin is no longer sufficiently protected against USP1-UAF1 deubiquitination. An interesting observation is that USP2 and USP7 have some activity towards D2_Ub_, but this activity is lost when the latter forms a complex with FANCI. This indicates that formation of a locked ID2_Ub_-DNA complex blocks D2_Ub_ from general DUB activity. However, USP1-UAF1 can deubiquitinate ID2_Ub_-DNA complexes and D2_Ub_ alone to the same degree. Hence, USP1-UAF1 is able to circumvent any protection formed from a locked ID2_Ub_-DNA complex. It is not until FANCI is also ubiquitinated (forming an I_Ub_D2_Ub_-DNA) that USP1-UAF1 activity towards D2_Ub_ is reduced. These data suggest that USP1-UAF1 may be capable of modulating the ID2_Ub_-DNA complex in order to gain access to FANCD2’s ubiquitin. Although the exact details by which this is achieved are not known, our recent work has shown that a FANCD2-binding sequence, located at USP1’s N-terminus, is required for ID2_Ub_-DNA deubiquitination [29]. Nevertheless, our work here shows that this ability of USP1-UAF1 is compromised when FANCI is also ubiquitinated. The effect of FANCI’s ubiquitination may be on rendering the doubly ubiquitinated ID2 complex unamenable to USP1-UAF1 modulation; for example by stabilising an ID2 conformational change induced by FANCD2 ubiquitination, or inducing further minor conformational changes that stabilize the complex. Our deubiquitination assay data with I_Ub_ ubiquitin mutants indicate that the ubiquitin on FANCI interacts with FANCD2, which is consistent with the recently reported I_Ub_D2_Ub_-DNA structure. Hence, it is possible that FANCI ubiquitination blocks USP1-UAF1 access to a FANCD2 binding site, which is required for cleaving the ubiquitin on FANCD2. In support for this, our previous work has shown that deletion of a FANCD2-binding sequence in USP1 similarly results in compromised USP1-UAF1 activity towards FANCI-ubiquitin-deficient ID2_Ub_-DNA complexes [29].

The formation of a USP1-UAF1 resistant I_Ub_D2_Ub_-DNA complex likely serves to increase the *in vivo* half-life of the di-monoubiquitinated ID2 complex. In that way a threshold event is created whereby once two ubiquitins are installed on the ID2 complex the latter is committed for its function. In contrast, the intermediate ID2_Ub_ complex is transient and rapidly deubiquitinated. Moreover, we observe that FANCD2 is deubiquitinated by USP1-UAF1 faster than FANCI in the IUbD2_Ub_ complex. This observation implies that an ordered deubiquitination mechanism may be in place, resulting in an IUbD2 complex intermediate. Previous work has shown that when DNA is removed from the reaction, di-monoubiquitinated ID2 is no longer resistant to deubiquitination by USP1-UAF1 [29,30], but It is currently not clear how DNA may protect I_Ub_D2_Ub_ from USP1-UAF1 deubiquitination. One possibility is that in the absence of DNA ubiquitinated FANCD2 dissociates from ubiquitinated FANCI, and in isolation both proteins are known to be susceptible to USP1-UAF1 activity [29]. However, the effect of DNA binding on the interaction between FANCI and FANCD2 in different ubiquitination states is unknown and will require description beyond 1:1 binding models. Nevertheless, removing DNA might be an effective way of achieving rapid ID2 deubiquitination and switching off the pathway. This could be achieved in a cellular context is by extraction of ubiquitinated ID2 complexes by the DVC1-p97 segregase [36] and subsequent deubiquitination by USP1-UAF1. Future work will be required to understand the sequence of events, DNA-dependence, and when other factors are involved in in I_Ub_D2_Ub_ deubiquitination.

Here we have uncovered distinct functions for two specific monoubiquitination events on FANCI and FANCD2 within the ID2 complex. We find that FANCD2 ubiquitination enhances binding of the ID2 complex to dsDNA and FANCI ubiquitination protects the complex from deubiquitination by USP1-UAF1. Combined, both events lead to a stable ID2 complex on dsDNA.

## Materials and Methods

### Cloning and mutagenesis of expression constructs

Constructs encoding for human FANCD2 having an N-terminal 3C-cleavable His_6_-tag (His_6_-3C-FANCD2), and human FANCI having N-terminal His_6_ and V5 tags separated by a TEV-cleavable site (His_6_-TEV-V5-FANCI), were described previously [26]. Related *FANCD2* and *FANCI* constructs encoding for a His_6_-3C-FLAG-tagged FANCD2 or His_6_-FANCI proteins were produced by site-directed mutagenesis. The R1285Q mutation was introduced into His_6_-FANCI by site-directed mutagenesis in the respective *FANCI* construct. Human *USP7* was ligated into a pFBDM vector by restriction cloning to encode for His_6_-3C-USP7. N-terminally His_6_-TEV-tagged USP1 (with G670A/G671A mutated auto-cleavage site [37]) and UAF1 were produced via sequential insertion of human *USP1* and *UAF1* cDNAs into an appropriate pFBDM vector by restriction cloning. All other constructs have been reported previously [25,26,29]. The coding regions of all constructs were verified by DNA sequencing.

### Protein expression and purification

All FANCI and FANCD2 constructs were expressed in *Sf21* insect cells and purified by Ni-NTA affinity chromatography, anion exchange, and gel filtration as previously described [26]. In some cases the His_6_-tag was removed from His_6_-3C-FANCD2 or His_6_-3C-FLAG-FANCD2 by 3C protease cleavage, prior to the gelfiltration step. Ubiquitinated FANCI and FANCD2 were produced and purified following *in vitro* reactions with Spy-3C-tagged ubiquitin, covalent linkage of ubiquitinated proteins with GST-tagged SpyCatcher, capture of resulting products on glutathione beads and subsequent cleavage of ubiquitinated proteins by 3C-protease treatment, as described previously [26]. For ubiquitinated His_6_-TEV-V5-FANCI and Flag-FANCD2 used in EMSAs, the steps including the reaction with Spy-3C-tagged ubiquitin and subsequent covalent linkage of ubiquitinated proteins with GST-SpyCatcher, were instead replaced by reactions with GST-3C-tagged ubiquitin. The 3C-protease treatment of ubiquitinated FANCD2 also resulted in removal of the six-histidine-tag from ubiquitinated FANCD2 constructs. After the final gel filtration step, purified proteins in GF buffer (20 mM Tris pH 8.0, 400 mM NaCl, 5% glycerol, 5 mM DTT or 0.5 mM TCEP) were cryo-cooled in liquid nitrogen in single use aliquots. The His_6_-FANCI R1285Q mutant was expressed and purified in the same way as the wild-type one.

Preparation of bacmids and protein expression in S*f*21 insect cells for USP1-UAF1 and USP7 was performed as previously described for FANCI and FANCD2 [26]. All steps following cell harvesting and prior to protein storage were performed at 4 °C. Cells were harvested ~72 hours following baculovirus infection. Cells were pelleted by centrifugation and cell pellets were resuspended in fresh ice-cold lysis buffer (50 mM Tris pH 8.0, 150 mM NaCl, 5% glycerol, 5 mM β-mercaptoethanol, 10 mM Imidazole, EDTA-free protease inhibitor cocktail (Pierce), 2 mM MgCl_2_, and Benzonase). Cells were lysed by sonication and clarified (32,000 x g for 45 min) before proteins were bound to Ni-NTA resin. Ni-NTA-bound His6-TEV tagged USP1-UAF1 complex and His6-3C tagged USP7 were further washed with 50 mM Tris pH 8.0, 150 mM NaCl, 5% glycerol, 1 mM TCEP, 10 mM Imidazole and then eluted into low salt buffer (50 mM Tris pH 8.0, 100 mM NaCl, 5% glycerol, 1 mM TCEP) containing 250 mM Imidazole. Anion exchange was performed for both USP1-UAF1 and USP7 by binding proteins to a ResourceQ (1 mL) column and eluting over a linear gradient (20 column volumes) of NaCl (100-1000 mM) in 20 mM Tris pH 8.0, 5% glycerol, 1 mM TCEP. Fractions containing USP1-UAF1 were subject to His-TEV protease overnight (ratio 1:10 protease to tagged protein). His_6_-TEV tagged USP1 and His_6_-TEV protease was bound to Ni-NTA resin, cleaved USP1-UAF1 complex was collected in the flow-through. In order to remove excess USP1, cleaved USP1-UAF1 was then concentrated and further purified using GL 10/300 Superdex 200 Increase column in 20 mM Tris pH 8.0, 150 mM NaCl, 5% glycerol, 5 mM DTT. Fractions from the peak containing the protein of interest were concentrated to ~5 mg/mL, and cryo-cooled in liquid nitrogen as single use aliquots. For His_6_-3C tagged USP7, protein was concentrated and subject to two rounds of gel filtration using a GL 10/300 Superdex 200 Increase column in 20 mM Tris pH 8.0, 150 mM NaCl, 5% glycerol, 5 mM DTT. The center of the asymmetric gel filtration peak for USP7 was collected from the first run and purified again on gel filtration, the final peak was symmetric and concentrated to ~ 5 mg/mL for flash freezing as single use aliquots in liquid nitrogen.

GST-USP2 was purified as previously described and the GST tag was retained [29]. Protein ubiquitination reagents (Uba1, Ube2T/Ube2Tv4, FANCL^109-375^) were prepared as described previously [26]. Protein concentrations were determined using absorbance at 280 nm and predicted extinction coefficients based on the protein sequences [38].

### DNA substrates

700IRDye-labelled dsDNA (ds32^F^) was obtained from annealing of two complementary 32-nucleotide-long HPLC purified 5′-end 700IRDye-labelled DNA ssDNA molecules. Non-labelled dsDNA (ds32 & ds50) was obtained from annealing of two PAGE purified complementary (32-nucleotide-long or 50-nucleotide long) ssDNA molecules. The 64-nulceotide long ssDNA (ss64) was similarly PAGE purified. The above dsDNAs were purchased from Integrated DNA Technologies in their final purified and/or annealed form, were subsequently resuspended at appropriate stock concentrations in distilled water, and stored at −20 °C until use. ssDNA was purchased from Sigma Aldrich, resuspended at appropriate stock concentrations in 10 mM Tris pH 8.0, 25 mM NaCl, 0.5 mM EDTA, and stored at −20 °C until use. Sequence details of all oligonucleotides used are provided in Table S1.

### Assessment of FANCI/FANCD2 ubiquitination or subsequent deubiquitination by SDS-PAGE, and Coomassie staining or western blotting

Non-ubiquitinated/deubiquitinated FANCI or FANCD2 proteins were distinguished from respective ubiquitinated products/substrates following SDS-PAGE on Novex 4-12 % Tris-glycine gels (ThermoFisher) and subsequent staining of the gels with InstantBlue Coomassie stain (Expedon). All samples loaded for SDS-PAGE (~300 ng of FANCD2 for Coomassie staining and ~100 ng for western blotting) were first diluted with reducing LDS buffer [consisting of NuPAGE 4x LDS buffer (ThermoFisher) and appropriate concentration of beta-mercaptoethanol] to 1x LDS and 100 mM beta-mercaptoethanol or 100 mM DTT final, and then heated for 2 min at 100° C. For assessing the progress of deubiquitination, ubiquitinated/deubiquitinated FANCI and FANCD2 were additionally visualised by western blotting. SDS-PAGE-separated proteins were transferred onto nitrocellulose membranes using an iBlot gel transfer device (Invitrogen) set at P0 (20 V 1min, 23 V 4 min, 25 V 2 min) and blocked with 5% milk PBS-T (0.05% tween 20) before incubation with 1:1000 rabbit anti-FANCD2 (sc-28194; Santa Cruz Biotechnology) and 1:100 mouse anti-FANCI (sc-271316; Santa Cruz Biotechnology) or anti-V5 (66007.1-Ig; ProteinTech) for 60 minutes at room temperature, or overnight at 4 °C. Membranes were washed extensively with PBS-T, incubated with secondary infrared-labelled antibodies (Li-Cor) for 90 minutes at room temperature and then washed extensively again with PBS-T. Bands were visualised on an Odyssey CLx (Li-Cor) using the 700- or 800-nm channel.

### Ubiquitination assays

Ubiquitination of isolated FANCI was performed using His_6_-FANCI or His_6_-FANCI^R1285Q^ (2 μM) as substrate, in the presence or absence of single-stranded DNA (ss64; 4 μM; Appendix Table S1). Reactions were conducted using His_6_-Uba1 (50 nM), His_6_-ubiquitin (5 μM), Ube2Tv4 (2 μM) and FANCL^109-375^ (2 μM) in a final reaction buffer of 49 mM Tris pH 8.0, 120 mM NaCl, 5% glycerol, 1 mM DTT, 2.5 mM MgCl_2_, 2.5 mM ATP. Ubiquitination of ID2 or I^R1285Q^D2 was performed in the presence of double-stranded DNA (ds50; Appendix Table S1). Reactions were prepared by dilution of all components into E3 reaction buffer (50 mM Tris pH 8.0, 75 mM NaCl, 5% glycerol, 5 mM MgCl_2_ and 2.5 mM ATP) to yield final concentrations of 4 μM FANCI (His_6_-FANCI or His_6_-FANCI^R1285Q^), 4 μM His_6_-3C-FANCD2, 16 μM ds50, 4 μM Ube2Tv4, 4 μM FANCL^109-375^, 100 nM His_6_-Uba1, and 8 μM ubiquitin. All reactions were performed at room temperature and stopped at indicated time-points by addition of reducing LDS buffer (1x final concentration).

### Deubiquitination assays

The ID2_Ub_-DNA and I_Ub_D2_Ub_-DNA substrates were reconstituted by mixing appropriate amount of D2_Ub_, His_6_-TEV-V5-FANCI or His_6_-TEV-V5-I_Ub_ and DNA (ds50; Appendix Table S1) to form a I/I_Ub_:D2_Ub_:DNA molar ratio of 1:1:4. D2_Ub_-DNA substrate were prepared in the same way, but with GF buffer substituting I/I_Ub_. The substrates were diluted in DUB buffer (50 mM Tris pH 8.0, 75 mM NaCl, 2 mM DTT, 5% glycerol) on ice to a concentration of 2 μM and incubated for at least 15 min. USP1-UAF1, USP7 or GST-USP2 were prepared at 2X concentrations in DUB buffer. To initiate reactions, 2X substrate was mixed 1:1 with 2X DUB in a 10 μL reaction volume and reactions were stopped at indicated time-points by addition of 10 μL of reducing 2X LDS buffer.

### DUB-step assays

His_6_-TEV-V5-FANCI (4 μM) was ubiquitinated using Ube2Tv4 (4 μM), FANCL^109-375^ (4 μM), His_6_-Uba1 (100 nM), wild-type or mutant ubiquitin (8 μM) and DNA (ds50; 16 μM). All reaction components (apart from ubiquitin) were first diluted using E3 reaction buffer (50 mM Tris pH 8.0, 75 mM NaCl, 5% glycerol, 5 mM MgCl_2_ and 2.5 mM ATP) and reactions subsequently initiated with addition of ubiquitin at room temperature. Reactions were arrested at indicated time-points by addition of 5 U/mL apyrase (New England Biolabs) and subsequent incubation on ice for 5 min. Ubiquitinated FANCI was subsequently mixed in a 1:1 ratio with purified D2_Ub_ (2 μM ID2, 8 μM DNA) and incubated for a further 15 min on ice. The reconstituted complexes, D2_Ub_, I_Ub_-DNA or I_Ub_D2_Ub_-DNA, were then subject to deubiquitination by USP1-UAF1 (final concentrations 100 nM USP1-UAF1, 1 μM substrate) in 10 μL reaction volumes for an indicated amount of time at room temperature. Reactions were stopped by addition of reducing LDS buffer (1x final concentration).

### Electro-mobility shift assays (EMSA)

Indicated FANCI and FANCD2 constructs were pooled at equimolar concentrations were serially diluted in GF buffer and each dilution was mixed with EMSA reaction buffer for final concentrations of 2 nM labelled DNA (ds32^F^; Appendix Table S1), 16 mM Tris-HCl, pH 8, 150 mM NaCl, 4.4 % glycerol, 0.07 mg/ml BSA, 7 mM DTT, 4 mM EDTA and ID2 concentrations ranging from 4 to 512 nM. Reactions (18 μl final) occurred on ice for 20 min, before addition of 2 μl of 10x Orange dye. 10 μl were then loaded on 4% polyacrylamide 0.5x TBE gels, that had been pre-run in 0.5x TBE buffer, and electrophoresis occurred at 135 Volts for 40-45 minutes. The gels were subsequently scanned in Li-Cor imaging system (Odyssey CLx) using the 700 nm laser. Percentage of DNA bound was determined by the ratio of protein-bound signal to total DNA signal per lane.

### Protein Induced Fluorescence Enhancement

For measurement of DNA binding, aliquots of FANCI (His_6_-FANCI or His_6_-TEV-V5-FANCI), His_6_-3C-FANCD2, or the ubiquitinated versions in GF buffer were thawed on ice, then mixed at a molar ratio of 1:1 to form each different ID2 complex. Samples were then exchanged into Fluorescence Buffer (20 mM Tris pH 8.0, 150 mM NaCl, 5% glycerol, 0.47 mg/mL BSA, 1 mM DTT) by 5-fold dilution with 20 mM Tris pH 8.0, 87.5 mM NaCl, 5% glycerol, 0.59 mg/mL BSA. Two-fold serial dilutions of exchanged protein were set up in PCR tubes with Fluorescence Buffer. Labelled DNA (ds32^F^), which was also diluted into Fluorescence Buffer, was mixed with each serial protein dilution to yield a final dsDNA concentration of 125 nM.

For measurement of FANCI-FANCD2 interaction, aliquots of FANCD2 (deriving from His_6_-3C-FANCD2 in which the His_6_-tag was cleaved by 3C protease) and His_6_-FANCI or His_6_-FANCI R1285Q were thawed on ice. Samples were then exchanged into Fluorescence Buffer by 5-fold dilution with 20 mM Tris pH 8.0, 87.5 mM NaCl, 5% glycerol, 0.59 mg/mL BSA. Two-fold serial dilutions of exchanged FANCD2 were set up in PCR tubes with Fluorescence Buffer. 120 nM exchanged His_6_-FANCI was labelled with 50 nM RED-tris-NTA dye (Nanotemper) in Fluorescence Buffer and added to the serial dilution to yield a final concentration of 60 nM His_6_-FANCI.

For measurement of DNA binding of FANCI ubiquitin mutants in complex with D2_Ub_, His_6_-TEV-V5-FANCI was ubiquitinated as per the DUB-step assays (using a reaction buffer of 10 mM Tris pH 8.5, 75 mM NaCl, 5% glycerol, 2.5 mM MgCl_2_ and 2.5 mM ATP) and terminated by addition of 5 U/mL apyrase. 15 μL of this reaction mix was added to 5 μL of His_6_-FANCD2_Ub_ (or GF buffer for the no D2_Ub_ control) to yield a final concentration of 3 μM I_Ub_D2_Ub_ (or I_Ub_) and a NaCl concentration of ~150 mM. Two-fold serial dilutions of this complex were set-up in matched buffer (including all the reaction components minus ubiquitin) and each was mixed in 1:1 ratio with labelled DNA (ds32^F^; 250 nM) in Fluorescence Buffer to yield a final dsDNA concentration of 125 nM.

Prior to fluorescence measurement samples were briefly centrifuged then transferred into premium capillaries (NanoTemper Technologies). Measurements were performed at 22 °C on a Monolith NT.115 instrument (NanoTemper Technologies) using the red channel. Laser power was set to 20% and 40% for DNA binding and FANCD2 binding, respectively.

### Fitting of binding data

Binding affinities and associated uncertainties were determined with GraphPad Prism by fitting a one-site binding model:

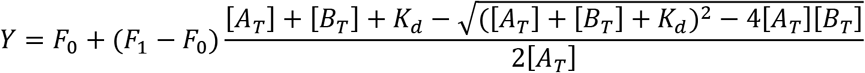

Where *Y* is either the fluorescence change (in the case of PIFE) or the percentage DNA-binding (in the case of EMSAs), *F*_0_ is baseline which is set to zero, *F*_1_ is the plateau, [*A_T_*] is the constant concentration of the fluorescent binding molecule, and [*B_T_*] is the varying concentration of the other binding molecule/complex. For binding curve of Figure 3F, the plateau *F*_1_ was set to the same *F*_1_ value calculated in Figure 1B. All datasets plotted together were assumed to have the same plateau i.e. a shared *F*_1_.

### CryoEM sample preparation

FANCD2_Ub_ and His_6_-TEV-V5-FANCI or His_6_-TEV-V5-FANCI_Ub_ aliquots were thawed and mixed at a molar ratio of 1:1 then exchanged into EM Buffer (20 mM Tris pH 8.0, 100 mM NaCl, 2 mM DTT) using a Zeba^™^ Spin 7K MWCO desalting column. For I_Ub_D2_Ub_-dsDNA, ds32 (Appendix Table S1) was then added at a molar ratio of 1:1:1 I_Ub_:D2_Ub_:DNA. The samples were diluted to 3 μM (I_Ub_D2_Ub_-dsDNA) or 1.5 μM (ID2_Ub_) of complex. 3.5 μL of sample was applied to glow discharged grids (C-Flat 2/2 or Quantifoil 2/2), blotted for 2.5-3.5 seconds, and cryo-cooled in liquid ethane using a Vitrobot operating at ~95% humidity at 4.5 °C.

### CryoEM data collection and image processing

For I_Ub_D2_Ub_-dsDNA, 1846 movies were collected on a Titan Krios (Thermo Fisher Scientific) equipped with a Falcon III detector operating in counting mode using EPU software (Thermo Fisher Scientific). Each movie was 60 frames and motion-corrected with dose-weighting using MotionCor2-1.1.0. For the ID2^Ub^ sample, 1146 and 916 movies were collected in two sessions on a 300 kV CRYOARM (JEOL) equipped with a DE64 detector operating in counting mode using SerialEM [39]. Each movie was 39 frames. For both datasets CTF correction was performed using gCTF [40]. Further data collection details are provided in Appendix Table S2.

Subsequence processing was performed using RELION 3.0 or 3.1 [41]. For the I_Ub_D2_Ub_-dsDNA dataset approximately 5,000 particles were manually picked and used to generate reference-free 2D class-averages. Selected class-averages were then used as templates to auto-pick particles. 350,878 particles were extracted and five rounds of reference-free 2D class averaging were used to remove poor particles. An initial model was generated using stochastic gradient descent followed by one round of 3D classification with 6 classes. Particles from the highest estimated resolution class (12,710) were then used in 3D refinement. A mask was then generated, and post-processing performed. For the ID2_Ub_ dataset particles 7,793 particles were manually picked and one round of reference-free 2D classification were performed to remove poor particles. An initial model was generated using stochastic gradient descent followed by one round of 3D classification with two classes. Particles from the best class (2,973) were then used in 3D refinement.

The mouse ID2 structure (PDB: 3S4W) [18] was fit as a rigid body into the final I_Ub_D2_Ub_-dsDNA map using UCSF Chimera [42]. Flexible fitting of Cα atoms was subsequently performed using iMODFIT [28] incorporating secondary structure constraints and using data to 17 Å. The fitted ID2 structure was then used to generate a map at 12 Å, which was subtracted from the experimental map using UCSF Chimera to identify the difference in density.

## Author contributions

M.L.R., C.A., K.L. and H.W. conceived this work; M.L.R., K.L., C.A. and V.K.C. purified proteins; M.L.R, K.L. and C.A. designed and executed experiments; M.L.R. performed PIFE and ubiquitination assays; K.L. performed EMSA experiments; C.A. and M.L.R. performed DUB-associated assays; M.C. prepared cryoEM grids; J.S. collected CRYOARM cryoEM data; M.L.R. performed cryoEM data analysis with guidance from L.S.; K.L. performed quantification and statistical analyses; K.L. made the figures with input from M.L.R; M.L.R, K.L. and C.A. wrote the manuscript; M.L.R, K.L. and C.A. edited the manuscript with contributions from all other authors; H.W. secured funding and supervised the project.

## Acknowledgements

We thank past and current members of the Walden laboratory for experimental suggestions, comments on the manuscript and their support. All constructs are available on request from the MRC Protein Phosphorylation and Ubiquitylation Unit reagents Web page (http://mrcppureagents.dundee.ac.uk) or from the corresponding author. We acknowledge Diamond Light Source for time on Krios IV under proposal EM16637 and thank Dr. Daniel Clare for collecting data. We acknowledge the Scottish Centre for Macromolecular Imaging (SCMI) for access to cryo-EM instrumentation, funded by the MRC (MC_PC_17135) and SFC (H17007). We thank Ms. June Southall (University of Glasgow) for assistance for the PIFE experiments. This work was supported by the EMBO Young Investigator Programme to H.W.; the European Research Council (ERC-2015-CoG-681582 ICL_Ub_ consolidator grant to H.W.

---

**Appendix Table S1.**
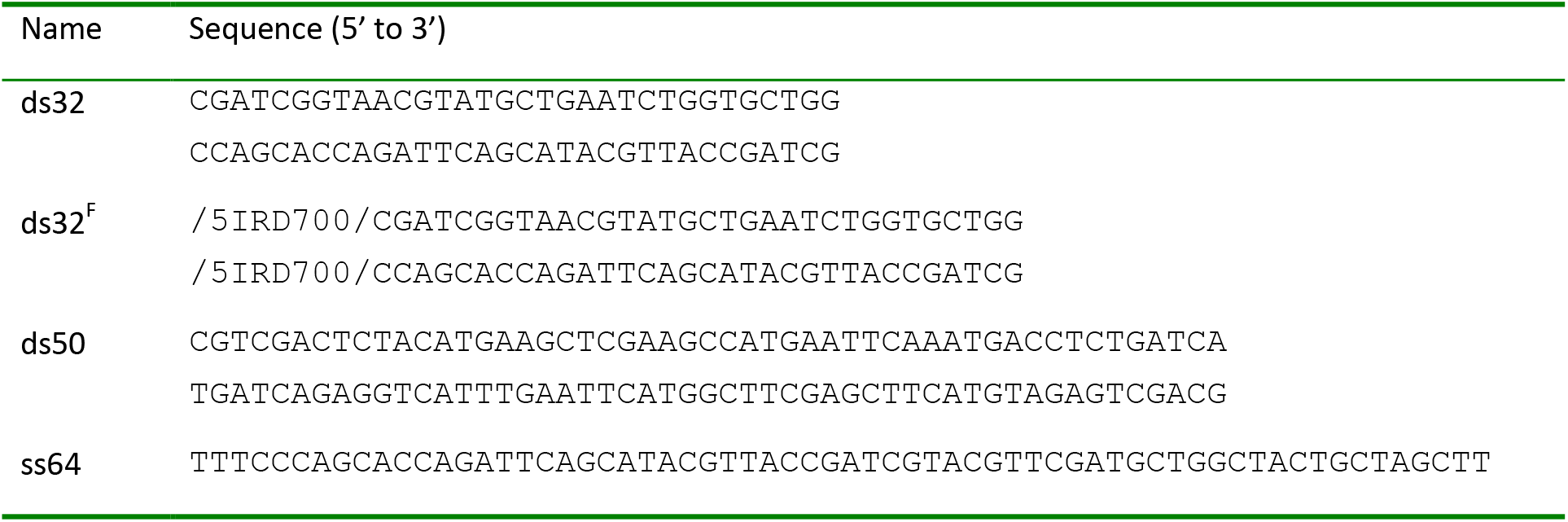
DNA substrates

**Appendix Table S2.** CryoEM data collection

|  | I<sub>Ub</sub>D2<sub>Ub</sub>-dsDNA | ID2<sub>Ub</sub> |
| --- | --- | --- |
| Microscope | Titan Krios | Cryoarm |
| Detector | Falcon III | DE64 |
| Magnification | 80,000x | 200,000x |
| Voltage (kV) | 300 | 300 |
| Electron Dose (e<sup>-</sup>/Å<sup>2</sup>) | 44.4 | 46.2 |
| Defocus range (μm) | 1.0-3.5 | 1.0-3.5 |
| Pixel Size (Å) | 1.085 | 0.598 |

**Figure EV1.**
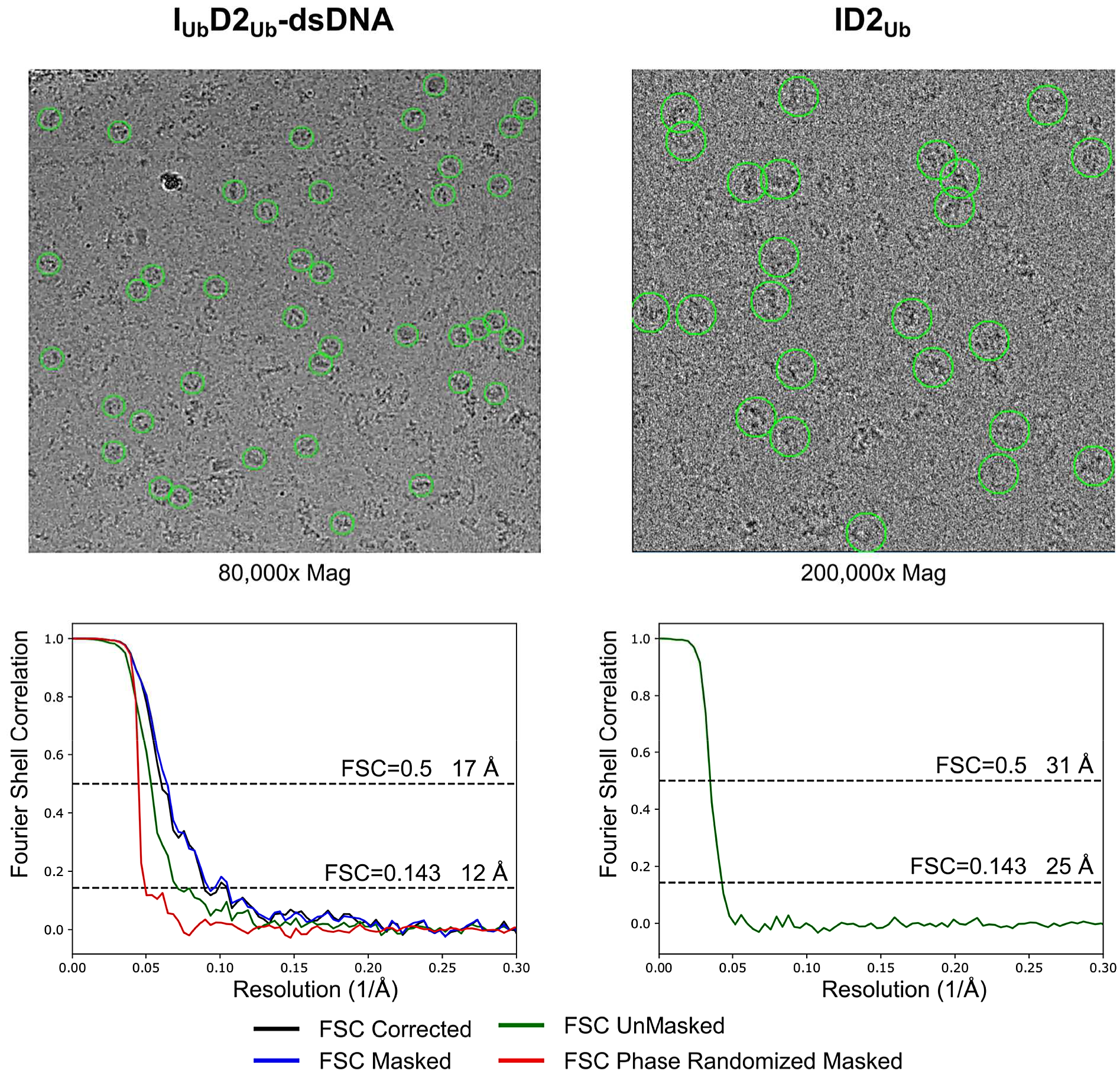
CryoEM data analysis for IUbD2Ub-dsDNA and ID2_Ub_ samples. *Top*: Selected micrographs with manually picked particles indicated with green circles. *Bottom*: Fourier Shell Correlation (FCS) curves for each dataset. For the IUbD2Ub-dsDNA dataset masking was performed with phase randomization to 23 Å and the indicated resolutions are for the corrected FSC curve.

**Figure EV2.**
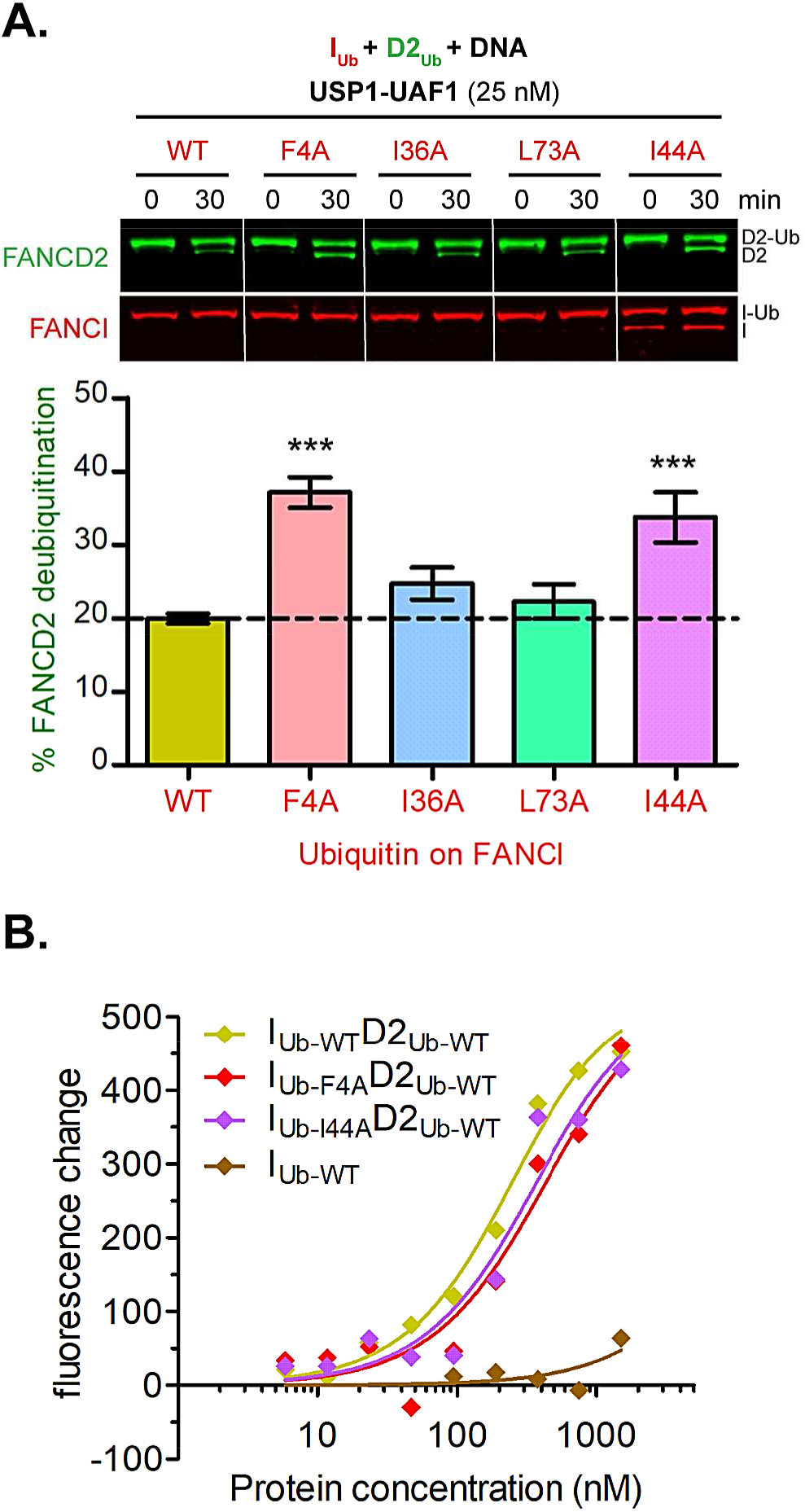
F4 and I44 in FANCI’s conjugated ubiquitin are required for efficient protection of FANCD2’s ubiquitin from USP1-UAF1 mediated deubiquitination. **A.** FANCI was first ubiquitinated *in vitro* in the presence of excess dsDNA, using wild-type ubiquitin (WT) or indicated ubiquitin-mutants. Ubiquitinated FANCI products were then mixed with ubiquitinated FANCD2 and resulting IUbD2Ub-DNA complexes were incubated with USP1-UAF1 (25 nM) for 30 minutes. Deubiquitination of FANCD2 and FANCI at 0 and 30 minutes was monitored by western blotting. The % FANCD2 deubiquitination over this period, calculated from FANCD2 blots deriving from three replicate DUB experiments, was plotted for each ubiquitin type (mean ± SD). Statistically significant changes compared to WT ubiquitin (one-way ANOVA test with Bonferroni correction) are indicated with asterisks. *** p<0.001. **B.** I_Ub_D2_Ub_ complexes with F4A or I44A ubiquitin mutants on FANCI can efficiently associate with dsDNA. PIFE DNA binding curves for I_Ub_D2_Ub_ complexes consisting of D2_Ub_ and I_Ub_ produced with either wild-type (WT) ubiquitin, or indicated ubiquitin mutants. IRD700-labelled dsDNA (at 125 nM) was incubated at increasing concentrations of corresponding IUbD2_Ub_ complexes or I_Ub_ control (ranging from 5.86 nM to 1.5 μM) and recorded fluorescence changes were plotted for each protein concentration along with the fit to a one-site binding model.

**Figure EV3.**
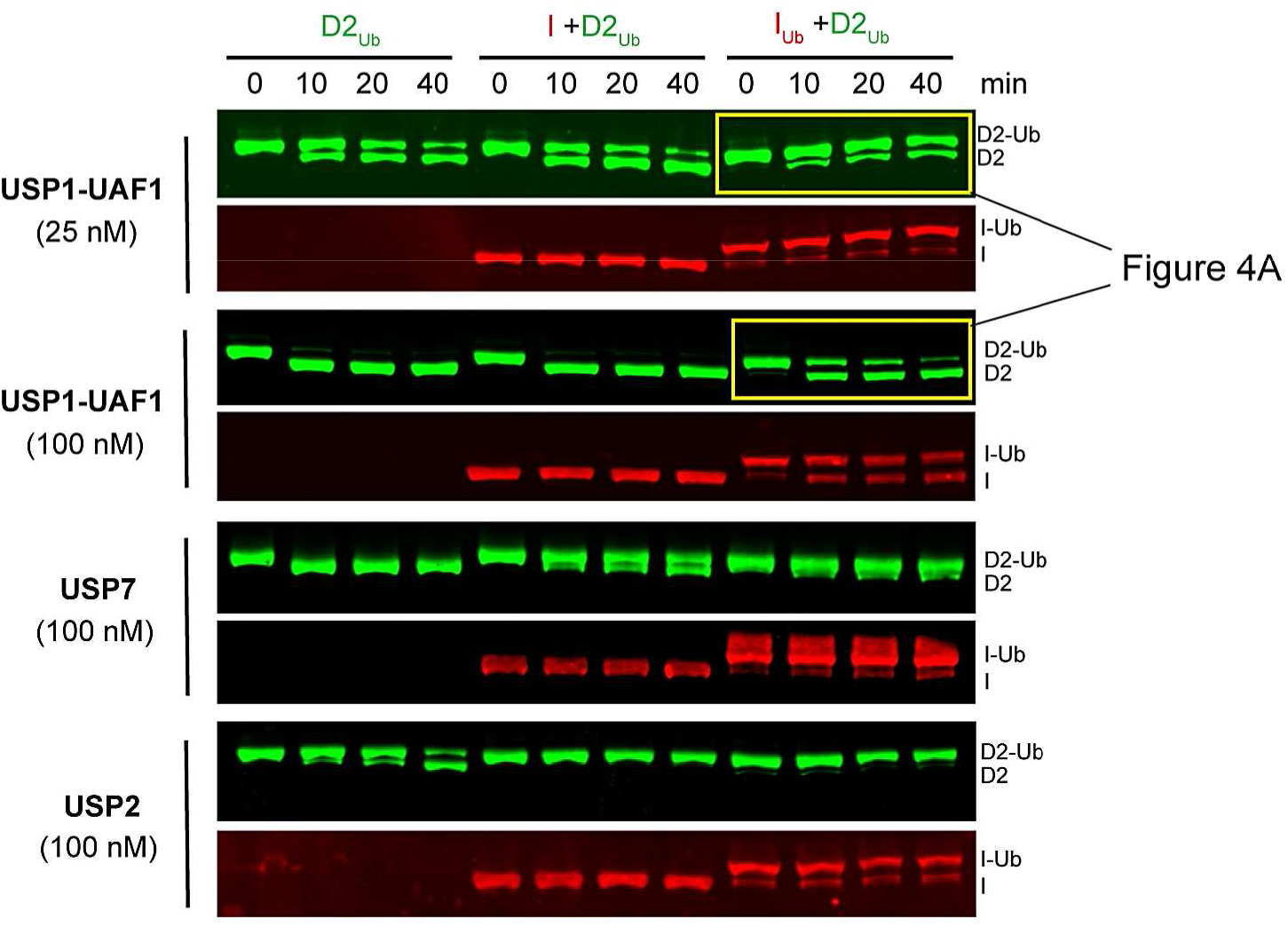
ID2_Ub_ complex is resistant to general DUB activity, whereas IUbD2_Ub_ is additionally resistant to USP1-UAF1 activity. Western blots of reactions in Figure 4A and 4B. Ubiquitinated His_6_-3C-FANCD2 (D2Ub) was mixed with a 50-nucleotide-long dsDNA and with either His_6_-TEV-V5-FANCI (I), ubiquitinated His_6_-TEV-V5-FANCI (IUb) or no protein; protein-DNA mixes were subsequently incubated with either USP1-UAF1 (at 25 nM or 100 nM), USP7 (100 nM) or USP2 (100 nM), for indicated time points. Deubiquitination of D2_Ub_ and I_Ub_ was assessed, following SDS-PAGE, by western blotting of transferred blots with specific FANCD2 and V5/FANCI antibodies.

